# Systematic literature review about educational interventions evaluated through entomological indices or practices to prevent the presence of or eliminate breeding sites of *Aedes*

**DOI:** 10.1101/2023.05.22.541679

**Authors:** Carola Soria, Walter Ricardo Almirón, Anna M. Stewart-Ibarra, Liliana Beatriz Crocco

## Abstract

**Background:** Community participation is a critical element in the management of *Aedes* breeding sites. Many educational interventions have been conducted to encourage prevention and elimination of breeding sites among different community actors, such as government-run programs for vector surveillance aimed at preventing and eliminating breeding sites at the household level within a community. Getting people involved in prevention and elimination of vector breeding sites in their communities requires communication and social mobilization strategies to promote and reinforce those prevention actions that, in turn, should be effective from the entomological standpoint.

**Methodology/Principal Findings:** Articles published in English, Spanish, and Portuguese, were reviewed to assess whether educational interventions targeting *Aedes* were effective in reducing entomological indicators or in improving practices to prevent the presence of or eliminate breeding sites. The most widely used indicators were the larval indices, and the practices to reduce/eliminate breeding sites. We found that using a community-based approach adapted to the eco-epidemiological and sociocultural scenarios explain the reduction of entomological indicators by educational interventions.

**Conclusions/Significance:** Those who design or implement educational interventions should strengthen the evaluation of those interventions using qualitative approaches that provide a more complete picture of the social context and barriers/facilitators to implementing vector control. Engaging school children in cross-sectorial collaboration involving the health and education spheres promotes the participation of the community in vector surveillance and reduces the risk of arboviral disease transmission.

**Author summary:** Dengue, Zika, and chikungunya are mosquito-borne diseases that represent a major global public health problem. These diseases are transmitted mainly through the bite of the *Aedes aegypti* vector mosquito and, to a lesser extent, *Ae. albopictus*. Getting people involved in prevention and elimination of mosquitos in their communities requires communication and social mobilization strategies to promote and reinforce those prevention actions that, in turn, should be effective from the entomological standpoint. The success of vector control programs has been demonstrated to lie in a comprehensive effort involving key community participation and intersectoral alliances. In addition, the participation of schoolchildren to mobilize their families in the prevention of breeding sites and the management of mosquito populations is recommended. In this article, we proposed to conduct a systematic review of scientific publications that evaluate the effects of educational interventions on *Aedes* through entomological indicators. As a result, we obtained only 26 articles that evaluated the efficacy of educational interventions in reducing vector populations out of 732 articles reviewed. The selected articles were published in both English and Spanish, and to a lesser extent in Portuguese, which highlights the importance of avoiding language bias in systematic reviews. As a conclusion of our work, we can mention that the interventions that incorporated the social context and the barriers/facilitators for the implementation of vector control were the most successful. In addition, we emphasize the importance of involving schoolchildren to promote community participation in vector surveillance.

## Introduction

Arboviral diseases, including dengue, yellow fever, chikungunya and Zika, are major public health problems in tropical and subtropical areas [1] and temperate regions where they are emerging [2, 3]. Half of the world’s population is estimated to be at risk of dengue infections. Every year, 100 and 400 million dengue infections occur [4, 5]. Dengue is transmitted by the *Aedes aegypti* mosquito vector and, to a lesser extent, *Ae. Albopictus* [5]; other mosquito species can be vectors in specific geographic contexts [6]. *Ae. aegypti* and *Ae. albopictus* can transmit other arboviruses, like Zika, Chikungunya, and Yellow Fever [7].

Preventing dengue and other arboviruses critically depends on controlling vectors to interrupt virus transmission. This strategy will likely remain a priority even when a vaccine becomes available [8, 9]. Successful vector management programs depends on comprehensive efforts to involve communities and cross-sectoral partners [10].

In this context, community education is crucial in promoting actions to eliminate vector breeding sites [11]. The World Health Organization (WHO) and the Pan American Health Organization (PAHO) [12] recommend, as a primary strategy, establishing a link with schools, with direct involvement of schoolchildren to mobilize their families in the prevention of breeding sites and management of mosquito populations [13].

Getting people involved in the prevention and elimination of vector breeding sites in their communities requires communication and social mobilization strategies to promote and reinforce prevention actions that, in turn, should be effective from the entomological standpoint [14]. In the case of dengue, the presence and abundance of vector populations are evaluated using entomological indicators, which are understood as proxies for dengue risk (presence of breeding sites, house index, container index, Breteau index, pupae per person index, and pupae per hectare index) [15].

Several articles have analyzed educational interventions aimed at promoting knowledge and prevention practices about dengue and other arboviral diseases; however, to date, no systematic reviews have explored the effect of those interventions on vector populations. The aims of the present work were: i) to conduct a systematic review of scientific publications that evaluate the effects of educational interventions on *Ae. aegypti* through entomological indicators and ii) to evaluate the effect of the interventions in reducing entomological indices and improving vector control practices.

## Method

A systematic literature review was conducted using the following databases: Medline (PubMed), Science Direct (Scientific Electronic Library Online), LILACS (Latin American and Caribbean Health Sciences Literature) and Redalyc (Network of Scientific Journals of Latin America and the Caribbean, Spain, and Portugal). For this, the terms “Dengue” and “*Aedes aegypti*” were combined with “education”, “schools” and “educational interventions” by the Boolean connector AND, and were used to search in the article title or abstract. The terms were adapted to the search modes of each database; thus, in the case of PubMed, the descriptors “education”, “schools” and “educational interventions ‘’ were replaced with the terms [MeSH] (Medical Subject Headings) “health education” and “education”.

### Eligibility criteria and search strategy

The search was restricted to the 2011-2021 period. The search terms used were both in Spanish and English to avoid introducing a language bias. In turn, no language restrictions were applied; therefore, articles published in other languages were not excluded. For an article to be included in the review, it had to: i) be published in indexed journals, ii) have free access to full text, and iii) be developed in formal/informal education contexts. Studies with an economic, sociological or epidemiological focus were not included.

Article selection for in-depth analysis was a two-stage process. During the first stage, two researchers identified and screened the titles and abstracts independently, following the eligibility criteria. The articles were incorporated into a predesigned spreadsheet with the items: database where they were included, year of publication, first author, journal, matches with the selection criteria and remarks on any observations that may arise. The search results of both researchers were compared and imported into Mendeley free software, and duplicates were removed. In the second evaluation stage, the articles were thoroughly read and those that evaluated the impact of education interventions through entomological indicators (house index, container index and Breteau index, or the presence and/or abundance of *Ae. aegypti* in any of its development stages) or prevention practices related to the elimination or control of breeding sites were included. The article selection process is presented in Figure 1.

**Figure 1.**
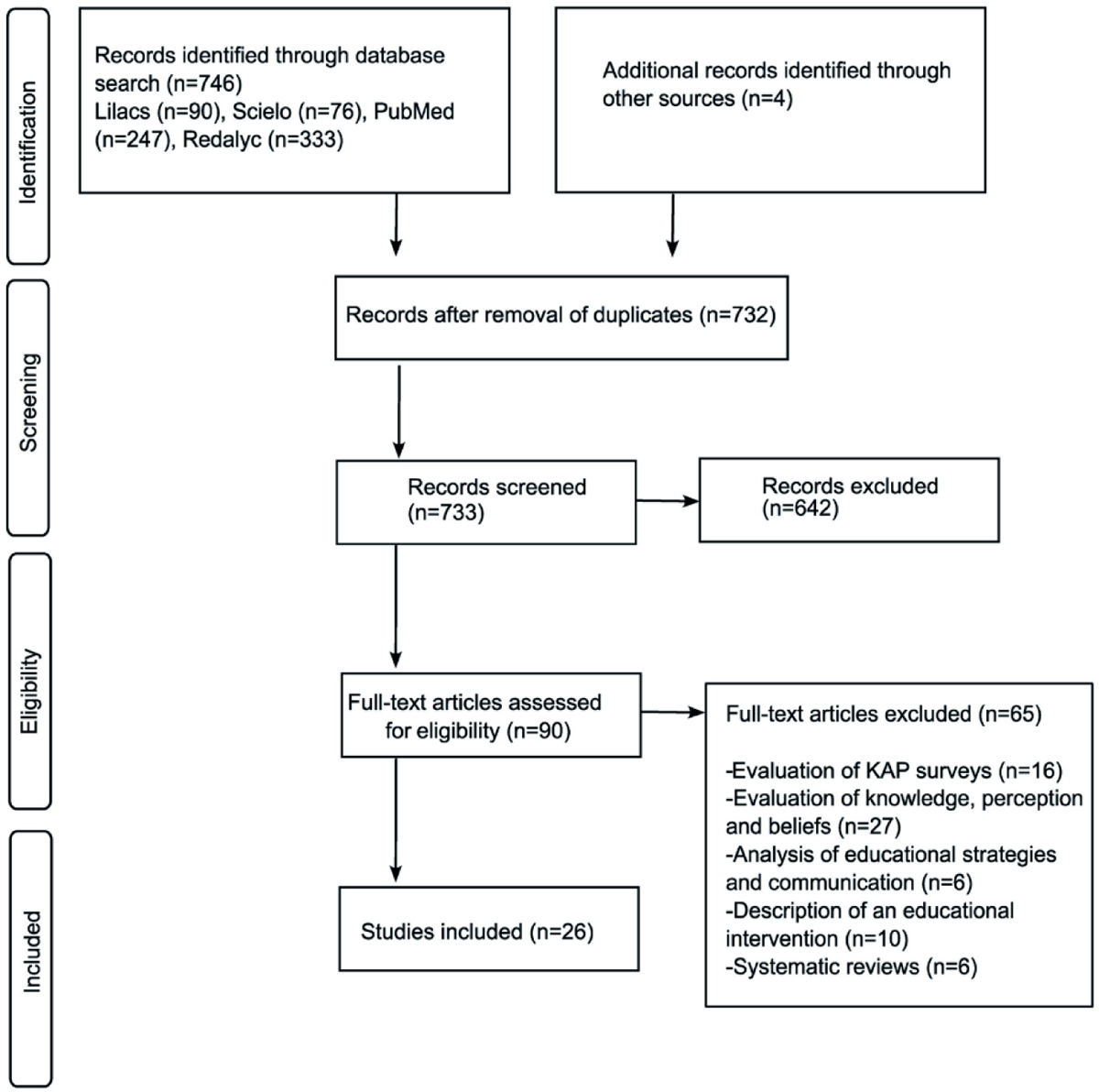
Process for the selection of published articles to be included in the systematic literature review.

Finally, the references cited in the selected articles were also reviewed to include other relevant articles that were not found during the database search. Any discrepancies during the first or second selection stages were resolved by mutual agreement between the reviewers. Quality of the analysis presented in the articles was not evaluated. In this sense, the risk of including articles with biased results or bias between articles was not assessed.

### Analysis

The selected articles were coded using the software Atlas Ti. The topics found during reading of the articles were summarized in descriptive topics. In a subsequent interpretation stage, the descriptive topics identified during this process were used to elaborate the criteria to select the articles. The articles were classified according to working aim, study area and target group of the educational intervention. The data presented in the works were not analyzed quantitatively.

## Results

Of all the articles searched, a small percentage (3.5%; 26/746) evaluated educational interventions using entomological indicators and most of them focused exclusively on *Aedes aegypti*. The interventions were conducted in South and Central America and in Asia, during years of high incidence of dengue and other arboviruses. No educational interventions evaluated by entomological indicators were implemented on temperate regions of the world. Community mobilization through educational interventions had a positive impact on the management of *Aedes* breeding sites.

### Description of the selected articles

The articles selected were published in Spanish, English and Portuguese. Most of the studies were conducted in South and Central America, mainly in Colombia, followed by South East Asia, and South Asia (Fig 2, Table 1). The authors used a total of 37 indicators to evaluate the educational interventions (Table 2).

**Figure 2.**
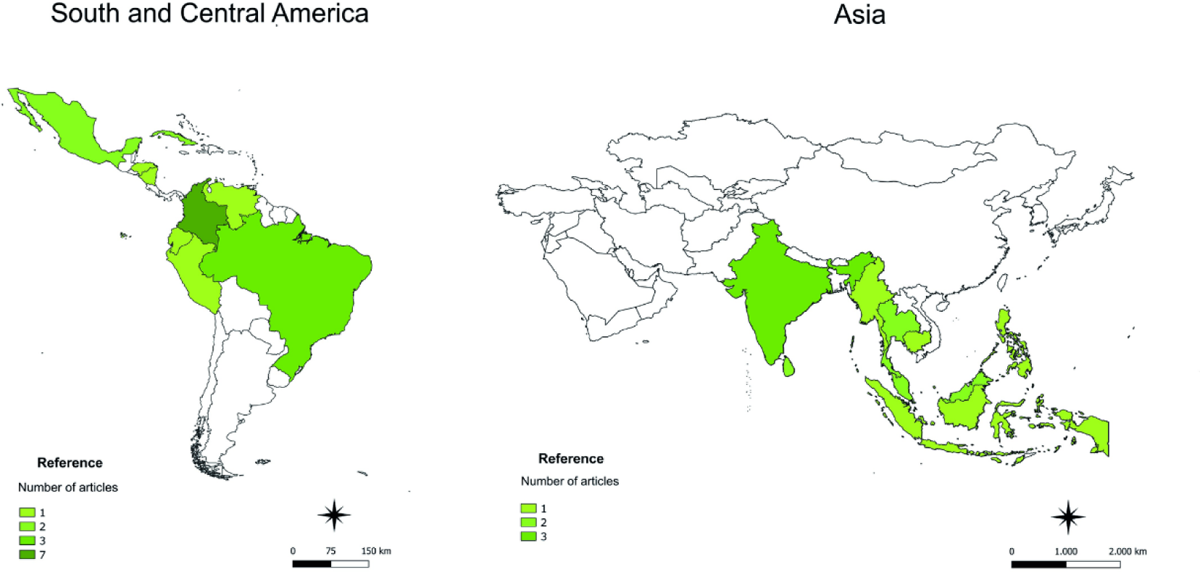
Distribution of articles evaluating educational interventions about *Aedes* spp. through knowledge, attitudes and practice (KAP) surveys and entomological indicators.

**Table 1.** Description of the articles selected for in-depth analysis

| Country | Number | Year of publication | Authors | Title | Target group |
| --- | --- | --- | --- | --- | --- |
| **South and Central America (18 articles)** |  |  |  |  |  |
| Colombia | 7 | 2011 | Restrepo et al. [35] | Aplicación y evaluación de materiales educativos para la prevención del dengue en una institución educativa de Medellín, Colombia. | Primary schoolchildren |
|  |  | 2013 | Carabalí et al. [21] | Difusión masiva de reportes situacionales sobre dengue: Efectos de la intervención en Guadalajara de Buga, Colombia. | Community |
|  |  | 2013 | Escobar Ramírez et al. [25] | Efectividad de la estrategia para la prevención del dengue en un barrio del municipio de Floridablanca 2011-2012. | Community |
|  |  | 2015 | Escudero Támara & Villareal Amaris [26] | Intervención educativa para el control del dengue en entornos familiares en una comunidad de Colombia. | Primary schoolchildren and community |
|  |  | 2016 | Overgaard et al. [33] | A cluster-randomized controlled trial to reduce diarrheal disease and dengue entomological risk factors in rural primary schools in Colombia. | Primary schoolchildren |
|  |  | 2017 | Villareal et al. [38] | Intervención educativa para control de *Aedes aegypti* en un grupo de familias colombianas: una experiencia exitosa. | Primary schoolchildren and community |
|  |  | 2018 | Jaramillo et al. [29] | Sostenibilidad en intervenciones para la prevención de dengue y diarrea en escuelas rurales de dos municipios de Colombia: evaluación de dos años post-proyecto. | Primary schoolchildren |
| Brazil | 3 | 2019 | Abel Mangueria et al. [16] | The prevention of arboviral diseases using mobile devices: a preliminary study of the attitudes and behaviour change produced by educational interventions. | University students |
|  |  | 2015 | Caprara et al. [20] | Entomological impact and social participation in dengue control: a cluster randomized trial in Fortaleza, Brazil. | Primary schoolchildren and community |
|  |  | 2013 | Cordeiro da Silva et al. [22] | Cooperação entre agentes de endemias e escolas na identificação e controle da dengue. | Primary schoolchildren, vector control agents and community |
| Cuba | 2 | 2012 | Zayas Vinent et al. [39] | La intersectorialidad en la prevención del dengue en un área de salud de Santiago de Cuba. | Community |
|  |  | 2016 | Criado Morales et al. [23] | Procesos de participación comunitaria en la prevención del dengue barrio ciudadela del fonce. | Community |
| Honduras | 1 | 2012 | Ávila Montes et al. [19] | Un programa escolar para el control del dengue en Honduras: del conocimiento a la práctica. | Primary schoolchildren and community |
| Mexico | 1 | 2014 | Torres et al. [37] | Conocimientos, actitudes y prácticas sobre el dengue en las escuelas primarias de Tapachula, Chiapas, México. | Primary school children |
| Nicaragua and México | 1 | 2015 | Andersson et al. [18] | Evidence based community mobilization for dengue prevention in Nicaragua and Mexico (Camino Verde, the Green Way): cluster randomized controlled trial. | Community |
| Peru | 1 | 2020 | Ortiz Agui et al. [32] | Estrategia comunicativa orientada a la reducción de la exposición a factores de riesgo de arbovirosis. | Primary schoolchildren |
| Venezuela | 1 | 2011 | García Guevara et al. [27] | Una mirada etnográfica del conocimiento del dengue desde la perspectiva de un grupo de escolares. Unidad educativa nacional bolivariana "Armando Zuloaga Blanco" Distrito Capital, Venezuela. 2009. | Secondary school student |
| Ecuador | 1 | 2015 | Mitchell-Foster et al. [41] | Integrating participatory community mobilization processes to improve dengue prevention: an eco-bio-social scaling up of local success in Machala, Ecuador | Primary schoolchildren and community |
| **Asia (8 articles)** |  |  |  |  |  |
| Malasia | 2 | 2013 | AbhiRami & Zuharah [17] | School-based health education for dengue control in Kelantan, Malaysia: Impact on knowledge, attitude and practice. | Secondary school students |
|  |  | 2020 | Isa et al. [28] | Mediational Effects of Self-Efficacy Dimensions in the Relationship between Knowledge of Dengue and Dengue Preventive Behaviour with Respect to Control of Dengue. | Community |
| India | 2 | 2012 | Arunachalam [40] | Community-based control of *Aedes aegypti* by adoption of eco-health methods in Chennai City, India. | Secondary school students and community |
|  |  | 2016 | Nivedita [31] | Knowledge, attitude, behaviour and practices (KABP) of the community and resultant IEC leading to behaviour change about dengue in Jodhpur City, Rajasthan. | Community |
| Camboya | 1 | 2020 | Echaubard et al. [24] | Fostering social innovation and building adaptive capacity for dengue control in Cambodia: a case study. | Primary schoolchildren and community |
| Tailandia | 1 | 2012 | Kittayapong et al. [30] | Application of eco-friendly tools and eco-bio-social strategies to control dengue vectors in urban and peri-urban settings in Thailand. | Community |
| Sri Lanka | 1 | 2019 | Radhika et al. [34] | Level of awareness of dengue disease among school children in Gampaha District, Sri Lanka, and effect of school-based health education programmes on improving knowledge and practices. | Secondary school students |
| India, Sri Lanka, Indonesia, Birmania, Filipinas, Tailandia | 1 | 2012 | Sommerfeld & Kroeger [36] | Eco-bio-social research on dengue in Asia: a multicountry study on ecosystem and community-based approaches for the control of dengue vectors in urban and peri-urban Asia. | Community |

**Table 2.** Indicators used in the reviewed articles to evaluate educational interventions about *Aedes aegypti* or *A. albopictus*.

| Indicators | Knowledge, attitudes and practice (KAP) survey | Entomological indicators of *Aedes aegypti* or *Aedes albopictus* |  |  |  |  |  |
| --- | --- | --- | --- | --- | --- | --- | --- |
|  |  | Presence of positive breeding sites (with larvae/pupae) | *Aedes* index * | HI, CI, BI† | Pupae per person index | Density of *Aedes aegypti* larvae and pupae per breeding site | Proportion of schools with adult females of *Aedes aegypti* |
| References | [16, 17, 28, 31, 32, 35, 37] | [30, 29, 38, 26, 22, 39, 19, 27, 34, 35] | [23, 24, 25] | [18, 19, 20, 21, 33, 35, 40, 41] | [18, 20, 21, 30, 33, 40, 41] | [36] | [33] |

\* Articles that evaluated the educational intervention through the *Aedes* index but did not define or specify how they calculated those indices. † References: HI: house index = number of houses with larvae of *Aedes* spp./total number of inspected houses x 100; CI: container index = number of positive containers with *Aedes* spp. larvae / total number of inspected containers with water x100; IB: Breteau index = Number of positive containers/ inspected houses x100

Of the educational interventions, two-thirds (65%) were performed in schools or involved the participation of schoolchildren [16, 17, 19, 20, 22, 24, 26, 27, 29, 32, 33, 34, 35, 37, 38, 40, 41]. One-third (35%) were focused on the community, with no participation of students [18, 21, 23, 25, 28, 30, 31, 36, 39]. The interventions that targeted the community encouraged home dwellers to perform activities to prevent or eliminate vectors in their homes through interventions designed and implemented by research groups [21, 23, 25, 28, 31] or vector control programs [18, 30, 36, 39].

Diverse education tools were used in the studies (Fig 3). Varona Delmonte et al. [42] defines educational tools as methods or activities used to teach a skill or concept. An important aspect to highlight is the heterogeneity among authors regarding the education tool used. Some studies [17, 21, 22, 23, 28, 31, 34, 35, 38] included a training event about concepts and prevention practices against dengue. In those events, the distribution of leaflets, talks, and screening of animated films were the prevailing features. These tools were used one to three times with the target group and were employed by research groups. Other studies [16, 18, 19, 20, 27, 29, 30, 32, 33, 36, 37, 39, 40, 41] applied theoretical-practical education tools. In particular, they conducted programs that lasted between 40 days and six years, which in general involved the coordination of different social actors in the design, implementation and evaluation of the educational intervention including teachers, researchers, community leaders, community health workers providing services on a voluntary basis (e.g., *brigadistas*), students, and members of health and environment departments. These studies mainly focused on home visits to detect breeding sites, focus groups, and community engagement in the design of the response to the problem.

**Figure 3.**
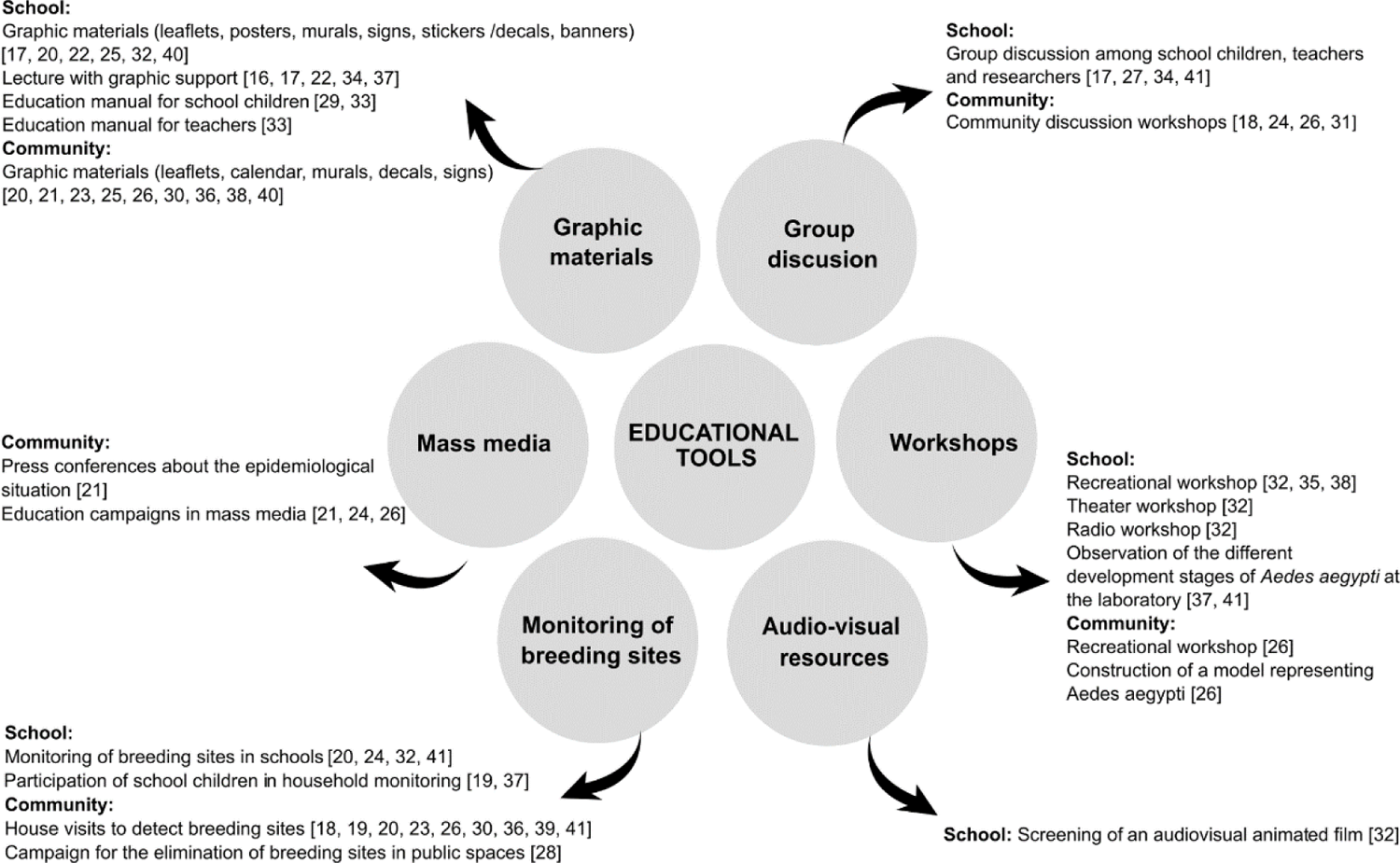
Educational instrument or tool used in the articles selected for the systematic review.

Most educational interventions focused on eliminating the presence of *Ae. aegypti.* Two studies included *Ae*. *albopictus*, [18, 34] and three studies referred to the vectors as *Aedes* mosquitoes [21, 26, 28].

### Evaluation of educational interventions through knowledge, attitudes, and practices (KAP) surveys

Some interventions, especially those related to the validation of education tools [16, 35, 37], communication strategies [32] or awareness-raising programs [17, 31], evaluated their effects on vector populations indirectly or through KAP surveys. The authors of those articles indicated that they educated people about dengue or mosquito vectors, which had a positive effect on the implementation of practices to prevent or eliminate adult vectors or larval habitats [16, 17, 31, 32, 35, 37]. They evaluated these practices through surveys including avoiding the presence of breeding sites [16], turning containers upside down, sealing containers hermetically, or washing containers to avoid water accumulation [31, 32], removing containers [35], and setting traps for adult mosquitoes [16].

Research groups conducted all of the KAP studies. Four were implemented in primary or secondary schools [17, 32, 35, 37], and one involved university students [16]. In contrast, only one was devoted to the community and consisted of surveying an adult in a house at the moment of the visit [31].

### Evaluation of educational interventions aimed at the community through entomological indicators

Of the interventions involving the general community, 90% used one of the entomological indicators [18, 21, 23, 25, 28, 30, 36, 39]. In general, the interventions resulted in a statistically significant reduction in entomological indicators, including the presence of breeding sites of *Ae. aegypti* larvae [30, 39], indices of *Ae. aegypti* larvae [18, 21, 23, 28, 41], pupal indices [21, 18, 41], and the density of larvae and pupae per breeding site [36]. Other indicators, such as qualitative and quantitative indicators of empowerment, collaboration, and community mobilization, were also used in some studies [18, 25, 36].

The educational interventions were mostly designed based on a diagnosis of community knowledge about the role of vectors in dengue transmission. Diverse educational tools (presented in detail in Fig 3) were implemented to evaluated the effects of educational interventions on entomological indicators.

Other indicators to measure the effectiveness of the educational intervention included: the identification of risk factors favoring the presence of breeding sites in the houses [23], changes in human behaviors relative to vector prevention [25], and the environmental risks that made implementation easier or more difficult [39]. Isa et al. [28] state that changes in preventive behaviors will not be effective unless the intervention also increases people’s confidence in their capacity to perform such behaviors. In turn, Carabalí et al. [21] state that a fundamental element to stimulate community participation is providing feedback on the impact of an educational intervention.

Some studies focused on evaluating the effect of educational interventions on vector control compared to traditional chemical control. Andersson et al. [21], Kittayapong et al. [30] and Mitchell-Foster et al. [41] presented evidence of the greater effectiveness of physical control of potential vector breeding sites coupled with educational interventions compared to the conventional chemical control of dengue. Similarly, Sommerfeld & Kroegel [36] reported the positive results of implementing interventions to reduce vector reproduction over five years in six Asian countries; interventions were based on education efforts, along with chemical, mechanical or biological treatments.

In some studies, key actors (community leaders, staff from the municipal program for vector control, health and education sector staff) were trained as *brigadistas*. Visited houses to show the dwellers the evidence of larval/pupa infestation in water containers, to assess general knowledge about dengue and to inform them about the mosquito life cycle and prevention measures [18, 30, 36, 41].

### Educational interventions with schoolchildren that were evaluated by entomological indicators

Of the 17 articles reporting the participation of schoolchildren, 71% used an entomological indicator to evaluate the effect of the intervention on vector populations. The authors concluded that the interventions were efficient in raising awareness and improving the attitudes of the school communities regarding the maintenance of a safe environment with a low level of vector infestation. However, some weaknesses were reported by Overgaard et al. [33], who highlighted that the favorable results –e.g., the knowledge acquired and practices for the prevention of breeding sites– observed during the intervention were not sustained after the intervention was finished.

Among the articles included in this section, educational interventions were implemented as part of the interventions for community-based ecosystem management, or Ecohealth approaches, to reduce entomological indices. The schoolchildren collaborated in community interventions, mainly as mobilizers and message disseminators [20, 24, 40] or in actions to eliminate mosquito breeding sites in their homes [41]. According to Echaubard et al. [24], implementing an Ecohealth strategy allowed them to achieve a cross-sector linkage among the community, teachers, heads of schools, and representatives of the Ministry of Education, who redefined the school curriculum to include dengue information. Other interventions proposed by research groups aimed at preventing breeding sites in houses by the training of schoolchildren [26, 38].

Using an ethnographic approach, García Guevara et al. [27] analyzed the presence of vectors by generating maps of the school with the distribution of the potential *Ae. aegypti* breeding sites. They evaluated students’ knowledge and compared it to the public health messages delivered via massive communication campaigns. These authors observed that students reproduced the campaign message with a reduced and decontextualized vision of prevention.

## Discussion

The results obtained in this review show that while there are numerous articles on educational interventions for the control of *Aedes*, few have evaluated these interventions using entomological indicators or assessments of vector prevention practices. It should be noted that 70% of the educational interventions were implemented in South and Central America, and 46% and 4% of the articles were published in Spanish and Portuguese, respectively. A strong feature of this systematic review is that it includes publications in languages other than English [43]. The high proportion of articles in Spanish points to the importance of including knowledge about dengue and other arboviral diseases published in local scientific journals in countries where *Ae. aegypti*-transmitted diseases pose a significant threat to community health. Most articles focused on *Ae. aegypti*; only two articles conducted in Malaysia and Sri Lanka [17, 34] considered *Ae. albopictus*. The lack of studies on *Ae. albopictus* is a key gap, given its potential role in arboviral transmission growing distribution of this mosquito species [44–46].

We found that educational interventions were performed mainly in schools or with the participation of schoolchildren, whereas the remaining interventions were aimed at the community. Among the latter, few studies clearly characterized the target community group or social actors. In this sense, we agree with Bang [47] in that there is a lack of consensus about the concept of *community* in disease prevention practices.

The articles selected in this review did not compare the efficacy of the different education tools applied in the interventions; instead, they evaluated the impact on various end points (improving knowledge and prevention practices or reducing entomological indicators). Al-Muhandis & Hunter [48] and Vázquez-Torres et al. [49] mention that no studies have been conducted to compare different education tools, presenting an exciting line of future research.

In the articles evaluated in this review, the most widely used tools to develop educational interventions were the monitoring of breeding sites in schools and homes and the distribution of graphic materials (flyers, manuals, posters, etc.). In some cases, the tools included games, theatre workshops, radio workshops, and an audiovisual presentation. The use of play-based strategies and the direct participation of students in controlling mosquitoes in their houses is recommended, since they were presented as the most attractive and effective education strategies to address dengue among children and adolescents in schools according to Diaz Gonzalez [50] in a systematic review on health education initiatives.

Regarding the evaluations of the interventions, one-third of the articles focused on schoolchildren used pre-test and post-test KAP surveys. Those questionnaires have been widely used to diagnose misconceptions or misinterpretations and to know the opinion of specific communities about a problem [51]. Several authors mention that educational interventions to address vector borne diseases (e.g., malaria, Chagas, dengue, chikungunya, and Zika) improved knowledge and induced positive changes in the practices to prevent and control vectors, as assessed by KAP surveys [52–56].

The most widely used entomological indicators were the presence of breeding sites and/or container, house and Breteau indices. This result is consistent with the review conducted by Ballenger-Browning & Elder about vector control interventions to reduce mosquito populations [57].

Several authors, like Ballenger-Browning & Elder [57], Heintzer et al. [58] and Erlanger et al. [59], discuss the use of larval indexes as entomological indicators to evaluate educational interventions and highlight that it was not possible to measure whether interventions that promote community participation in vector control were able to effectively reduce dengue transmission. These authors suggest that serological surveillance should be a necessary component of all dengue interventions and a standard entomological index should be used to compare interventions. In addition, entomological indices estimated through larva collection are imperfect indicators of disease transmission risk [60, 61]. Andersson et al. [18] was the only evidence found that incorporated – in addition to the reduction of entomological indicators - serological analysis as an indicator of the effectiveness of the educational intervention. But the results of the community serological analysis are not yet available.

Therefore, we cannot ascertain whether educational strategies assessed via entomological indicators successfully reduced virus transmission. Active surveillance studies and serological surveys would be needed to measure changes in disease transmission pre and post intervention. Instead, entomological indicators provide information to evaluate whether people adopt practices to reduce or eliminate vectors, which is the aim of educational interventions.

Some articles performed an in-depth analysis of the relationship between educational intervention and practices to achieve source reduction/elimination of breeding sites. The authors explain that incorporating actions to prevent breeding sites requires that people have confidence in the benefits of those actions to their health and that they receive feedback on the results of the interventions they participate in. Those who design or implement educational interventions should strengthen the evaluation of those interventions using qualitative approaches that provide a more complete picture of the social context and barriers/facilitators to implementing vector control.

Another group of articles conducted interventions from an eco-bio-social approach, involving different social actors. The results show that combining the education strategy with other vector control strategies contributed to the reduction of both larval indices and pupal densities. Andersson et al. [18] and Kittayapong et al. [30] mention that educational interventions framed as community-based Ecohealth strategies are more efficient in reducing vector populations and transmission than conventional chemical control. These findings are in agreement with results of Espinoza-Gómez [62]. Educational interventions made within the eco-bio-social frame and that involved community engagement were the most efficient vector control strategies at the global level [48, 49, 59, 62–64].

Several studies focused on school- or student-centered interventions to address a range of vector-borne diseases [65, 68]. The programs consisted of play-based activities to engage the students’ families in interventions to prevent the presence of or eliminate breeding sites in their houses. The main indicators evaluated were the reduction of the presence of breeding sites, house, container and Breteau indices, and to a lesser extent, pupal density. These results are encouraging since schools are essential in gathering community members, and students can act as a crucial link to transmit educational interventions [69].

Through a systematic review, Vázquez-Torres et al. [49] observed that the participation of students improves their knowledge about and application of dengue prevention measures, which had a significant effect on the reduction of entomological indicators in the school. The articles included in the present review agree with Vázquez-Torres et al. [49] on the importance of health education to improve knowledge and prevention practices regarding arboviral diseases, especially the role of school teachers in the application of those practices.

This systematic review intends to contribute information about the positive impact of conducting educational interventions on a globally important health problem. For that purpose, we consider that having indicators of the efficacy of interventions helps to improve the design of the intervention.

The terms used in the literature search were selected with the aim to incorporate all the articles with information about the impact of educational interventions on vector prevention practices or entomological indicators. However, a weakness of our study is that some articles that do provide information about the impact of educational interventions in the text may have been overlooked because the descriptors we used were not included in the titles or abstracts. Finally, this systematic review did not aim to evaluate the quality of the educational interventions implemented or the method used and the subsequent published data.

In conclusion, those designing or implementing educational interventions should strengthen the evaluation of such interventions using qualitative approaches that provide a comprehensive picture of the social context and barriers/facilitators to vector control implementation. Future evaluations of the relationship between community mobilization and prevention of arboviruses could include entomological indicators that provide information to assess whether people adopt practices to reduce or eliminate vectors, along with active surveillance studies and serological surveys to measure changes in disease transmission before and after the educational intervention. From a health education policy perspective, engaging school children in cross-sectorial collaborations that link the health and education spheres is essential, thereby promote community participation in vector surveillance.

## Acknowledgments

We thank Jorgelina Brasca for the grammar and style revision of the language.

